# *Fezf2* transient expression via modRNA with concurrent SIRT1 inhibition enhances differentiation of cortical subcerebral / corticospinal neuron identity from mES cells

**DOI:** 10.1101/2020.08.13.242230

**Authors:** Cameron Sadegh, Wataru Ebina, Anthony C. Arvanites, Lance S. Davidow, Lee L. Rubin, Jeffrey D. Macklis

## Abstract

During late embryonic development of the cerebral cortex, the major class of cortical output neurons termed subcerebral projection neurons (SCPN; including the predominant population of corticospinal neurons, CSN) and the class of interhemispheric callosal projection neurons (CPN) initially express overlapping molecular controls that later undergo subtype-specific refinements. Such molecular refinements are largely absent in heterogeneous, maturation-stalled, neocortical-like neurons (termed “cortical” here) spontaneously generated by established embryonic stem cell (ES) and induced pluripotent stem cell (iPSC) differentiation. Building on recently identified central molecular controls over SCPN development, we used a combination of synthetic modified mRNA (modRNA) for *Fezf2*, the central transcription factor controlling SCPN specification, and small molecule screening to investigate whether distinct chromatin modifiers might complement *Fezf2* functions to promote SCPN-specific differentiation by mouse ES (mES)-derived cortical-like neurons. We find that the inhibition of a specific histone deacetylase, Sirtuin 1 (SIRT1), enhances refinement of SCPN subtype molecular identity by both mES-derived cortical-like neurons and primary dissociated E12.5 mouse cortical neurons. *In vivo*, we identify that SIRT1 is specifically expressed by CPN, but not SCPN, during late embryonic and postnatal differentiation. Together, these data indicate that SIRT1 has neuronal subtype-specific expression in the mouse cortex *in vivo*, and its inhibition enhances subtype-specific differentiation of highly clinically relevant SCPN / CSN cortical neurons *in vitro*.

## Introduction

Subcerebral projection neurons (SCPN) are the broad population of cerebral cortex (cortical) neurons that connect and provide high-level descending control via axonal projections from the neocortex (termed “cortical” here) to distal targets in the brainstem (midbrain, hindbrain) and spinal cord (Molyneaux *et al*., 2007; Woodworth *et al*., 2012; Custo Greig *et al*., 2013). The large subtype of SCPN providing descending motor control to the spinal cord (direct or via sensory feedback) are termed corticospinal neurons (CSN), a term often considered to also include cortical neurons projecting to brainstem targets. SCPN are the brain neurons that degenerate in ALS and related motor neuron diseases, and whose injury (in particular, to CSN) is responsible for the loss of voluntary motor function in spinal cord injury (Sances et al., *Nat Neurosci.* 2016). Early and defining molecular features of SCPN include high-level expression of FEZF2 and CTIP2, required for the specification and control of SCPN molecular, cellular, and anatomical identity (Molyneaux *et al*., 2005; Arlotta *et al*., 2005; Chen *et al*., 2005; Ozdinler and Macklis, 2006; Lai *et al*., 2008; Chen *et al*., 2008; Shim *et al*., 2012; Woodworth *et al*., 2012; Greig *et al*., 2013; Woodworth *et al*., 2016).

Midway through corticogenesis, post-mitotic SCPN identity is initially masked by transient co-expression of regulators of interhemispheric callosal projection neuron (CPN) development, including SATB2 (Alcamo *et al*., 2008; Britanova *et al*., 2008; Azim *et al*., 2009; Woodworth *et al*., 2012; Sadegh and Macklis, 2014; Leone *et al*., 2015). At later stages of maturation, SCPN discontinue expression of SATB2, and further resolve into diverse subpopulations of FEZF2- and CTIP2-expressing projection neurons with cortical area- and target-specific molecular identities, properties, and functional circuit connectivity (Woodworth *et al*., 2012; Custo Greig *et al*., 2013; Woodworth *et al*., 2016; Greig *et al*., 2016; Galazo *et al*., 2016).

Multiple epigenetic factors support the post-mitotic identity refinement of contrasting cortical neuron subtypes, such as SCPN and CPN, and enable their maturation (Kishi and Macklis, 2004; MacDonald and Roskams, 2008, 2009; Kishi *et al*., 2012; Yip *et al*., 2012). For example, SATB2 is a matrix-attachment region (MAR) binding protein (Britanova *et al*., 2005; Gyorgy *et al*., 2008; Alcamo *et al*., 2008; Britanova *et al*., 2008) that can mediate long-range interactions of enhancer sites with promoters (Yasui *et al*., 2002; Cai *et al*., 2003; Dobreva *et al*., 2003), and, together with SKI, can recruit the nucleosome remodeling and histone deacetylase (NuRD) complex (Baranek *et al*., 2012). Moreover, the transcription factors CTIP2 and CTIP1 (BCL11B, BCL11A; Leid *et al*., 2004), which are differentially expressed with subtype-specificity in the cortex, and which regulate the precision of SCPN and CPN differentiation (Arlotta *et al*., 2005; Lai *et al*., 2008; Tomassy *et al*., 2010; Woodworth *et al*., 2016; Greig *et al*., 2016), have also been demonstrated to interact with both the NuRD complex (Topark-Ngarm *et al*., 2006; Cismasiu *et al*., 2005) and SIRT1 (Senawong *et al*., 2003; Senawong *et al*., 2005) to mediate chromatin remodeling in cells outside of the brain. Together, these reports suggest that chromatin remodeling might contribute to post-mitotic refinement of cortical projection neuron subtypes, particularly in refinement of CTIP2-expressing SCPN from SATB2-/CTIP1-expressing CPN.

ES/iPSC-based models of cortical differentiation are emerging as useful tools to investigate roles of chromatin modifications in cortical development (Tiberi *et al*., 2012; Juliandi *et al*., 2012). While protocols for directing cortical differentiation from ES/iPSC cells in systems ranging from monolayer cultures to organoids have succeeded in replicating some of the molecular characteristics of cortical development (Eiraku *et al*., 2008; Gaspard *et al*., 2008; Michelsen *et al.,* 2015; Lancaster *et al*., 2013, Arlotta, 2018; Pasca, 2018), the mature refinement of cortical subtypes is incomplete with these protocols; immature neurons become “stalled” at an mid-embryonic developmental stage (Sadegh and Macklis, 2014). These data suggest that ES-derived cortical cells are unlikely to have a sufficiently permissive molecular context for the precise refinement of SCPN identity. *In vivo*, *Fezf2* mis-expression in multiple embryonic and early postnatal forebrain progenitor and early post-mitotic neuron populations can redirect their differentiation to SCPN-like identities, suggesting a strategy to potentially circumvent this problem (Molyneaux *et al*., 2005; Lai *et al*., 2008; Chen *et al*., 2008; Rouaux and Arlotta, 2010; Rouaux and Arlotta, 2013; De la Rossa *et al*., 2013). However, *Fezf2* is regulated by multiple cofactors, including Sox family transcription factors (Lai *et al*., 2008; Azim *et al*., 2009; Shim *et al*., 2012), and, in the absence of a forebrain-specific molecular context at early stages of ES cell differentiation, *Fezf2* mis-expression by ES cells does not drive SCPN molecular identity (Wang *et al*., 2011; Miskinyte *et al*., 2018; Sadegh, unpublished data).

ES/iPS cell-derived models of neocortical differentiation necessarily bypass precisely orchestrated, spatiotemporal mechanisms of embryonic differentiation (as demonstrated in the analysis of intermediate states of ES cell-derived spinal motor neurons; Briggs *et al*., 2017), suggesting that, in contrast to established spinal motor neuron differentiation, additional subtype-specific epigenetic modulation might be required for optimal in vitro generation of diverse neocortical neuron subtypes.

We hypothesized that alteration of the chromatin landscape within incompletely specified mES-derived cortical progenitors might promote a permissive molecular context for *Fezf2*-directed SCPN subtype refinement. To identify candidate chromatin remodeling enzymes, we conducted a high-content screen of mouse mES-derived cortical cells using a library of small molecules that modulate known epigenetic enzymes. Mature SCPN refinement was assessed by measuring changes in the ratio of positive (CTIP2) and negative (SATB2, CTIP1) markers of SCPN differentiation. This strategy emphasizes the utility of multiple exclusionary markers to delineate SCPN-specific differentiation among mES-derived cortical progenitors, in contrast to the approach of evaluating for multiple positive markers that are often expressed in immature SCPN-like neurons (Sadegh and Macklis, 2014).

From this screen, we identify the histone deacetylase Sirtuin1 (SIRT1) as an effective repressor of *Fezf2*-mediated SCPN molecular refinement. Small molecule inhibitors of SIRT1 (*e.g.*, EX-527, CHIC-35) enhance *Fezf2*-induced molecular maturation of SCPN by maintaining CTIP2 expression and reducing SATB2 and CTIP1 expression by both mES-derived cortical-like neurons and primary dissociated mouse cortical neurons *in vitro*. We also identify differential refinement of SIRT1 expression in late embryonic cortical neuron subtypes *in vivo*: elevated SIRT1 expression by CPN, and diminished expression by SCPN. Together, these data identify chromatin remodeling as an important mechanism of cortical subtype refinement both *in vivo* and for *in vitro* directed differentiation, and identify a route to enhanced subtype-specific differentiation of developmentally and clinically important cortical neurons from pluripotent stem cells.

## Results

### ModRNA provides dose- and time-dependent protein expression in mES-derived cells

Synthetic modified mRNA (modRNA) enables precision over gene dosage and timing by multiple cell types (Warren *et al*., 2010). Because modRNA does not integrate into the genome, and has a limited duration of expression (∼2 days), modRNA enables transient gene expression without manipulating the genomes of ES-derived cells. We tested the feasibility of modRNA transfection in feeder-free E14Tg2a mES cells undergoing an established monolayer protocol of differentiation that generates heterogeneous, maturation-limited, neocortical-like neurons (Gaspard *et al*., 2008; Gaspard *et al*., 2009; Sadegh and Macklis, 2014).

In agreement with the prior literature, we find a dose-dependent intensity of GFP expression after transfection of mES-derived cells with *GFP* modRNA. At the peak of pallial-like differentiation at day 14, modRNA-induced GFP expression peaks between 12-24 hours, with a sharp reduction of expression by 48 hrs (**Figure 1A,B**). We also find that modRNA transfection is not biased to a specific neural population; modRNA broadly transfects NESTIN-expressing neural progenitors, TuJ1-expressing immature neurons, and other cells (**Figure S1**). There is no appreciable change in cell density between conditions, consistent with previously published work (Warren *et al*., 2010). Importantly, the timing and duration of modRNA expression matches the known kinetics for other proteins and transcription factors (Mandal and Rossi, 2013). These data indicate that modRNA transfection enables dose- and time-dependent gene expression in mES-derived cells, including progenitors and neurons.

**Figure 1.**
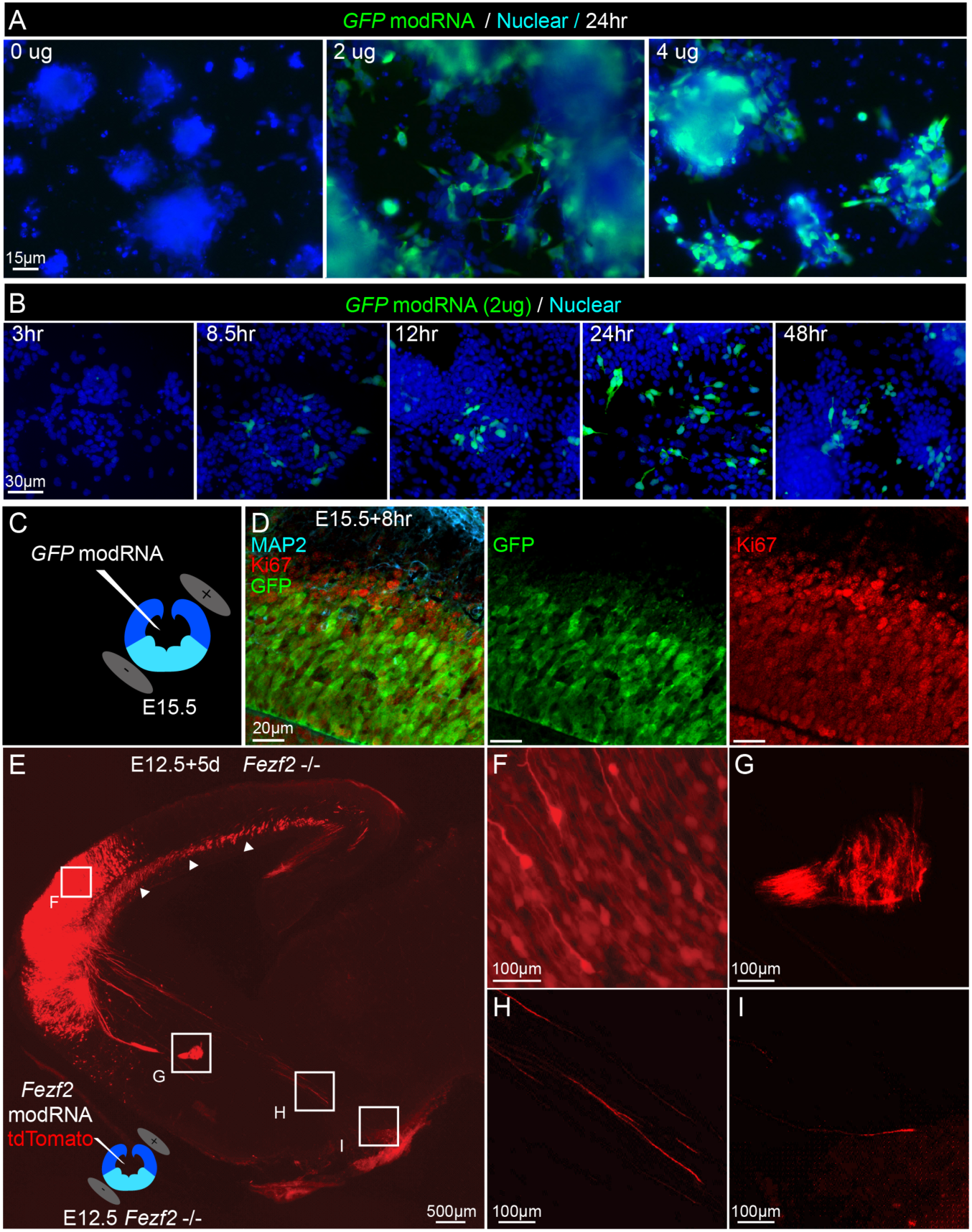
ModRNA enables temporally controlled, dose-dependent protein expression in cultured mES-derived populations, and *in vivo*. (A) *GFP* modRNA expression is dose-dependent. Native GFP expression is detected in 25-50% of cells after 24hrs; the intensity of expression increases over the range of 0, 2, 4 ug modRNA transfection. (B) *GFP* modRNA expression is time-dependent over 48hrs. (C) *GFP* modRNA is injected into the lateral ventricles (coronal cross-section of mouse embryo forebrain is shown), and is directionally electroporated into the dorsal forebrain, or pallium (positive paddle above dark blue colored tissue). (D) Following *GFP* modRNA *in utero* electroporation, GFP (green) is expressed after eight hours, and is restricted to Ki67-expressing progenitors (red) of the pallium, not in more superficially located post-mitotic neurons, which express MAP2 (blue). (E) Sagittal section of an E17.5 *Fezf2*-null mouse following E12.5 *in utero* electroporation of *Fezf2* modRNA (0.8 ug/uL) and *tdTomato* plasmid (1 ug/uL) shows targeted tdTomato fluorescence appropriately restricted to the rostral pallium. (F) Higher magnification image showing tdTomato expression in the cortical plate. (G) tdTomato+ axons project across the cerebral commissures, including anterior commissure (depicted) and corpus callosum in (arrowheads in E). (H) Several tdTomato+ axons project caudal to the thalamus, indicating partial genetic rescue of E12.5 SCPN, which do not exist at all in *Fezf2* null mice and can only be generated by the *in utero* rescue (Molyneaux *et al*., 2005). (I) A smaller subset of these tdTomato+ axons reach the cerebral peduncle (N = 3), confirming that they are SCPN.

### Transient Fezf2 expression rescues SCPN fate specification in Fezf2^-^/- mice *in vivo*

Prior to using *Fezf2* encoded modRNA for transient expression in mES-derived neurons, we first tested the functionality of *Fezf2* modRNA in mice *in vivo*, using an established *Fezf2* null experimental rescue paradigm. In *Fezf2* null mice, SCPN do not develop, and their progenitors are re-specified to CPN (Molyneaux *et al*., 2005; Chen *et al*., 2005). However, delivering *Fezf2* by *in utero* plasmid electroporation in *Fezf2* null mice at E12.5 (when FEZF2 would normally be expressed at a high level) can rescue SCPN specification and their projections to the distal hindbrain (Azim, 2009). Moreover, at E13.5, E15.5, and later ages, *Fezf2* mis-expression by plasmid electroporation can redirect CPN to acquire most critical features of SCPN identity (Molyneaux *et al*., 2005; Chen *et al*., 2008; Rouaux and Arlotta, 2010; Rouaux and Arlotta, 2013; De la Rossa *et al*., 2013).

We hypothesized that, if transient *Fezf2* expression delivered by *in utero* electroporation of a single dose of *Fezf2* modRNA in Fezf2 null mice can rescue SCPN differentiation, then a similar approach to modRNA delivery might effectively direct differentiation of ES-derived pallial-like progenitors *in vitro*. The extent of *in utero* GFP modRNA electroporation into the pallium (**Figure 1C**) is largely limited to the mitotic (Ki67-expressing) ventricular zone (**Figure 1D**). In *Fezf2* null mice at E12.5, we find that *in utero* electroporation of *Fezf2* modRNA (with tdTomato plasmid for long-term visualization of axons; **Figure 1E,F,G**) rescues a subset of SCPN that project beyond the thalamus to the cerebral peduncle (**Figure 1H,I**), comparable to similarly timed plasmid-mediated *Fezf2* expression using the same, robust *in utero* electroporation platform (Molyneaux *et al*., 2005; Arlotta *et al*., 2005; Chen *et al*., 2005; Chen *et al*., 2008; Shim *et al.,* 2012). In contrast, *in utero* electroporation of *Fezf2* modRNA does not induce SCPN specification at E15.5 (data not shown) when progenitors are no longer producing SCPN. These data suggest that a single, transient dose of *Fezf2* is sufficient to rescue *Fezf2*-null SCPN at E12.5, and that this relatively small dose of *Fezf2* modRNA is only functional within the permissive molecular context of E12.5 neocortical progenitors.

### Transient *Fezf2* expression alone does not significantly promote SCPN differentiation by mES-derived neocortical-like neurons

We next asked whether *Fezf2* modRNA alone can induce SCPN-specific differentiation by mES-derived neocortical-like cells (Gaspard *et al*., 2008; Gaspard *et al*., 2009; Sadegh and Macklis, 2014). Because these previously established protocols of monolayer differentiation generate limited quantities of mES-derived neocortical neurons, we used randomized, automated imaging (at 20x magnification, on approximately 40 fields per well; ∼5,000 cells), to count sufficient numbers of neocortical-like neurons for these analyses. A high threshold for positive antibody labeling was manually established because populations of mES-derived neurons express a continuum of transcription factor labeling intensities, in striking contrast to populations of primary dissociated E15.5 mouse neocortical neurons, which typically display distinct trimodal labeling (negative, low expression, high expression; see Methods for details). Normally, induction of CTIP2 expression occurs within 48 hrs of *Fezf2* plasmid expression. However, 48 hrs after Fezf2 modRNA transfection in mES-derived neocortical cells at *in vitro* day 18, the total numbers of either CTIP2- or SATB2-expressing neocortical neurons are not increased (**Figure S2A**). This result is not surprising given the heterogeneity and immaturity of neocortical-like neurons in this established protocol of ES cell culture (Sadegh and Macklis, 2014).

We hypothesized that, rather than broadly promoting CTIP2 expression in non-neocortical-like neurons, a transient dose of *Fezf2* modRNA might promote SCPN subtype-specific refinement by only a smaller subset of neocortical-like neurons, perhaps those already “poised” to differentiate further into corticofugal neurons. We developed a metric for delineating SCPN identity refinement in this subset of neocortical-like neurons by quantifying the ratio of neurons that have matured and only express CTIP2 to neurons that remain immature and have overlapping expression of CTIP2 and SATB2. Even by this more nuanced metric of SCPN identify refinement, we find that *Fezf2* modRNA expression in mES-derived neocortical cells does not, by itself, significantly increase SCPN subtype differentiation (**Figure S2A**,**B;** although there is a trend toward increased SCPN). These data indicate that a single, transient dose of *Fezf2* expression by mES-derived neurons is not sufficient to refine SCPN identity given the inappropriate molecular context of heterogeneous and maturation-stalled mES-derived neocortical-like neurons (Sadegh and Macklis, 2014), suggesting that additional, potentially complementary manipulations are needed to direct SCPN differentiation.

### Small molecule screening of mES-derived neocortical-like neurons identifies SIRT1

We next asked whether remodeling the chromatin landscape might enable a higher proportion of neocortical-like neurons to respond to *Fezf2*-mediated SCPN subtype refinement. To address this question, we designed an approach combining small molecule screening with transient *Fezf2* induction: 1) directed mES cell differentiation to day 14 neocortical-like progenitors, 2) addition of a small molecule library and incubation for four days to precondition the cells to a more permissive epigenetic landscape, 3) *Fezf2* modRNA transfection and incubation for two days to direct SCPN subtype refinement, and 4) immunohistochemical assessment using multiple exclusionary markers to evaluate the extent of SCPN identity refinement (**Figure S3A**). We designed a custom library of eighty small molecules modulating known epigenetic enzymes, with targets including histone deacetylases, methyltransferases, and kinases (**Figure S3B**). Using automated confocal imaging, cell segmentation, and threshold analyses, we quantified the expression of both CTIP2 and SATB2 by individual neurons (**Figure S3C,D**).

We used multiple selection criteria to identify leading candidates. In the first assay, we found that multiple Sirtuin modulators can either enhance or diminish *Fezf2*-mediated subtype refinement, as indicated by our metric of SCPN identity refinement, which is the ratio of maturing CTIP2^+^/SATB2^-^ neurons to relatively immature CTIP2^+^/SATB2^+^ double-positive neurons (**Figure 2A**). Focusing on small molecules that might enhance *Fezf2*-mediated refinement of CTIP2^+^/SATB2^-^ expression, we then asked which small molecules globally increase the total number of CTIP2 expressing neurons, relative to *Fezf2* modRNA induction alone (**Figure 2B**). In a third level assay, we tested a smaller group of leading candidate small molecules for their ability to either maintain or decrease the total number of SATB2 expressing neurons compared with *Fezf2* induction alone (**Figure 2C**). Optimized by these stringent criteria and given the prevalence of candidates independently targeting the same Sirtuin pathway, we chose the SIRT1 inhibitor EX-527 as a leading candidate to enhance *Fezf2*-mediated SCPN differentiation. As an internal control, we compared the activity of EX-527 to other Sirtuin inhibitors and activators. Non-specific Sirtuin inhibitors (nicotinamide, forskolin, and tenovin-6) do not increase SCPN refinement. Reinforcing these results, the SIRT1-specific *activator* (CAY10591) displays antagonism to SCPN refinement.

**Figure 2.**
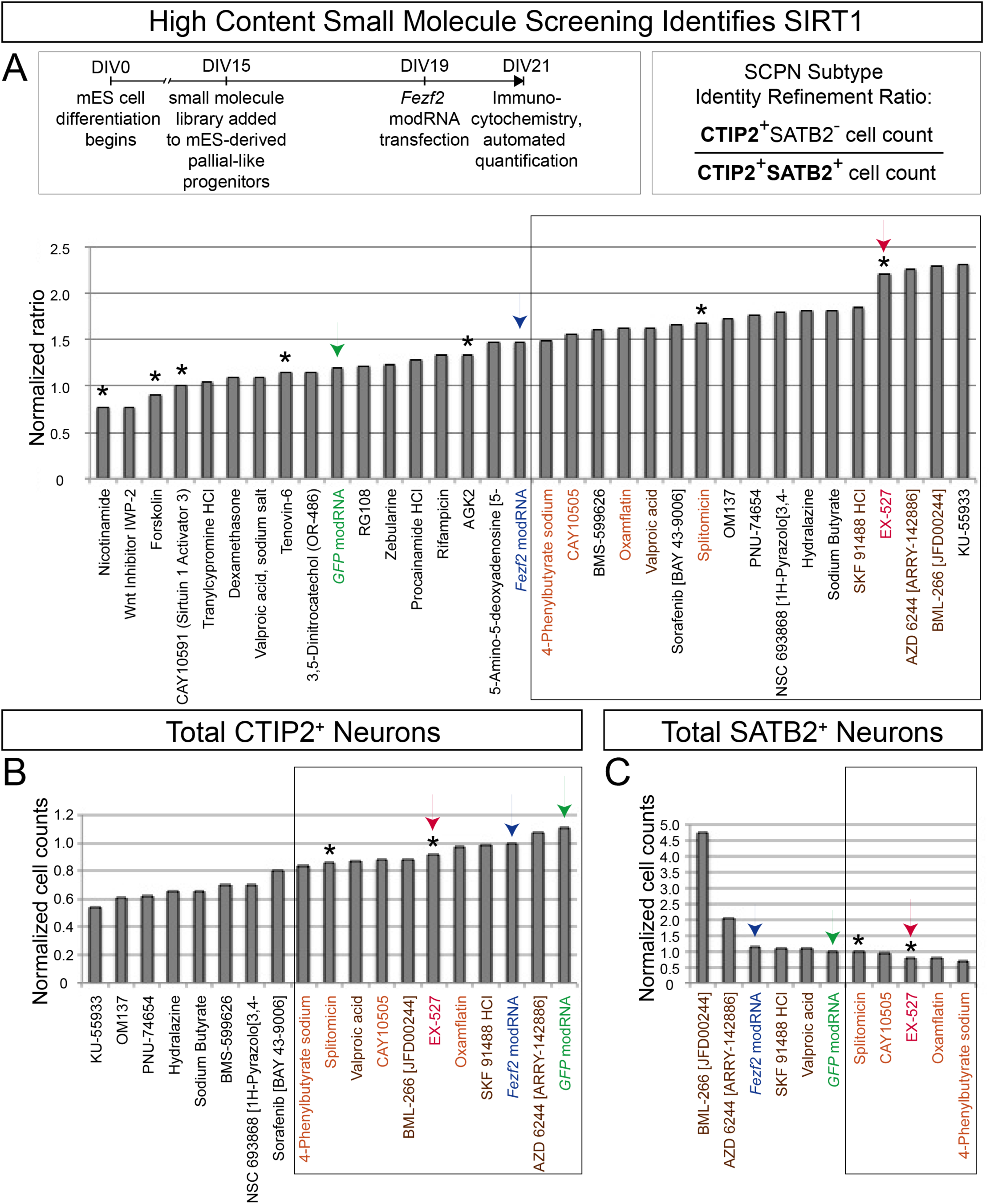
High-content small molecule screen of SCPN/CSN molecular refinement by mES-derived neurons identifies candidate small molecule regulators, including the SIRT1 inhibitor EX-527. (A) Each well of a 96-well plate containing ES-derived neurons at 15 days *in vitro* (DIV) was incubated with a distinct small molecule from the library (Figure S3B) for 96hrs, followed by transfection with *GFP* or *Fezf2* modRNA, incubated for an additional 48hrs, and then analyzed at 21 DIV. Distinct small molecules enhanced, inhibited, or did not substantially alter *Fezf2*-mediated SCPN subtype refinement, as indicated by the ratio of CTIP2(+)/SATB2(-) neurons to CTIP2(+)/SATB2(+) neurons. The ratios are normalized to the untreated condition. (B) Some candidate small molecules, from the box marked in (A), increase the number of CTIP2-expressing neurons relative to *Fezf2* induction alone. The ratios are normalized to the Fezf2 modRNA condition. (C) A subset of candidate small molecules, from the box marked in (B), decrease the number of SATB2-expressing neurons relative to *Fezf2* induction alone; these include the SIRT1 inhibitor EX-527. Asterisks indicate small molecules that modify HDAC Class III (Sirtuin). Black arrowheads indicate *GFP* and *Fezf2* transfection conditions, for comparison. The red arrowhead indicates EX-527, a relatively specific SIRT1 inhibitor. The ratios are normalized to the GFP modRNA condition. Data from this first-level screen experiment (N=1) represent approximately 1,000 cells per condition, from 40 randomly sampled fields at 20x magnification.

### SIRT1 inhibition refines primary dissociated E12.5 neuron SCPN subtype identity

We next asked whether SIRT1 inhibition also promotes SCPN subtype distinction by primary neocortical neurons. In these experiments, *Fezf2* was not induced with modRNA, because *Fezf2* is already highly expressed by primary SCPN progenitors. Dissociated E12.5 neocortical neurons were treated with small molecule inhibitors of SIRT1 for six days. We again identified EX-527, and an even more specific SIRT1 inhibitor, CHIC-35, as potent enhancers of CTIP2^+^/SATB2^-^ subtype identity refinement (**Figure 3A**). CHIC-35 is highly SIRT1-specific, with a binding site within the SIRT1 catalytic cleft that blocks substrate binding (Napper *et al*., 2005; Zhao *et al*., 2013). Compared to non-specific inhibitors, both EX-527 and CHIC-35 show selective enhancement of SCPN molecular refinement (**Figure 3A**). Notably, there was no observable change in the density of cultured cells following small molecule application, which was confirmed by automated cell segmentation analysis demonstrating stable cell counts, sizes, and fluorescence in the imaging wells across conditions.

**Figure 3.**
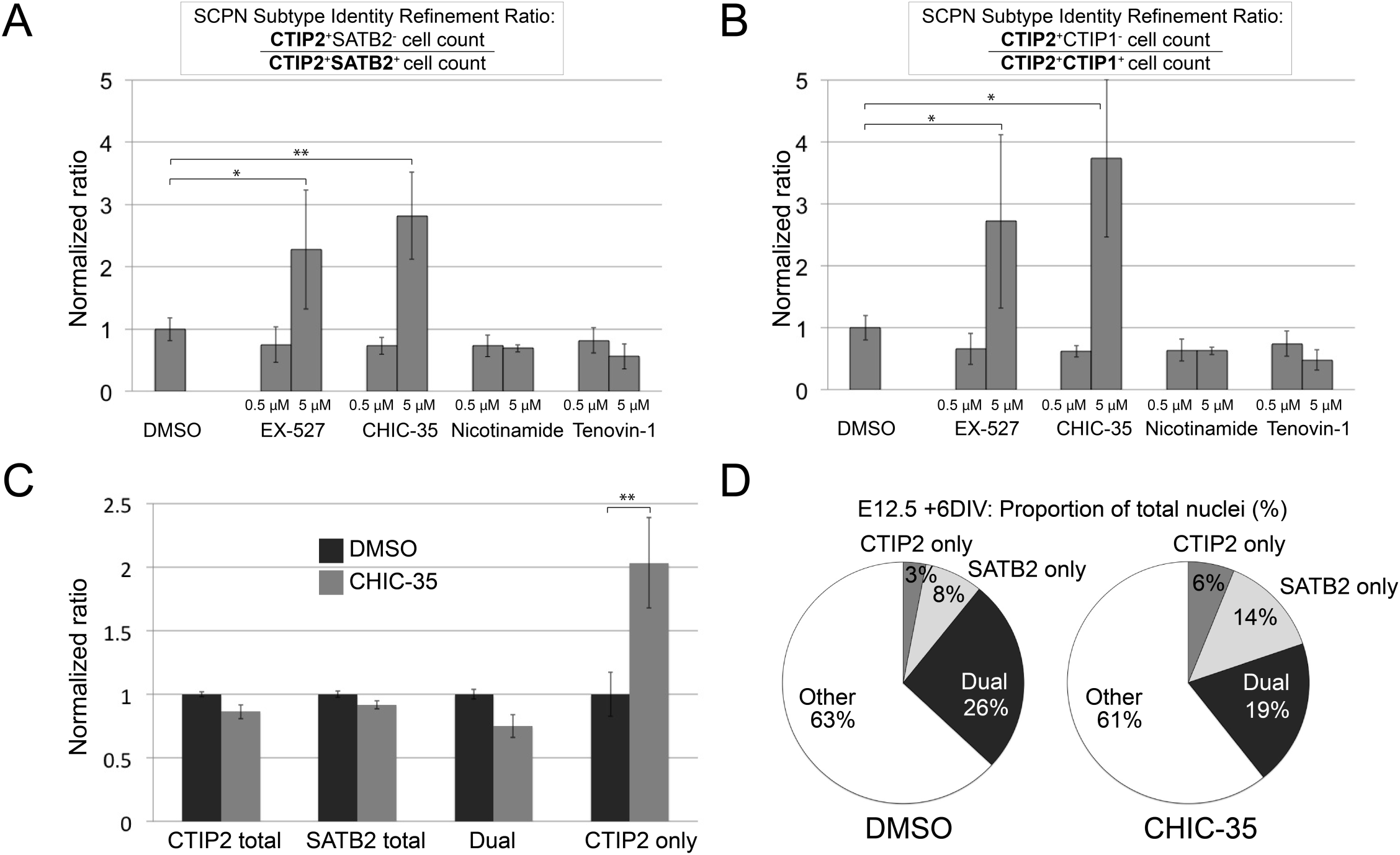
SIRT1 inhibition in dissociated E12.5 neocortical neurons enhances SCPN/CSN subtype refinement, increasing the number of CTIP2-expressing neurons at the expense of CTIP2/SATB2 dual expressing neurons. (A) EX-527 and a more specific SIRT1 inhibitor, CHIC-35, increase CTIP2/SATB2 subtype distinction relative to DMSO-only controls (0.5 μM and 5 μM). (B) EX-527 and CHIC-35 also trend toward CTIP2/CTIP1 subtype distinction. (C) Importantly, while the proportions of total CTIP2- and total SATB2-expressing neurons do not change relative to DMSO control following CHIC-35 SIRT1 inhibition, the relative proportion of CTIP2/SATB2 dual expressing neurons decreases. In important contrast, the relative proportion of more fully distinguished CTIP2(+)/SATB2(-) neurons (reflecting SCPN/CSN) increases. (D) Pie chart schematics show the relative proportions of total nuclei of CTIP2+-only neurons, SATB2+-only neurons, CTIP2+/SATB2+ dual expressing neurons, and unlabeled cells derived from E12.5 neocortical cells in CHIC-35 SIRT1 inhibition treatment conditions versus DMSO control. Data are presented as mean +/- s.e.m. (N=3; >10,000 nuclei screened per condition from 60 randomly sampled fields at 20x magnification) *P < 0.05; **P < 0.01 (unpaired t-test).

To further investigate whether SIRT1 inhibition broadly regulates SCPN subtype identity, rather than potentially only downregulating SATB2 expression, we asked whether other subtype-specific refinements occur. CTIP1 is a transcription factor that regulates both subtype- and area-specific identity (Woodworth *et al*., 2016; Greig *et al*., 2016). Despite its close homology to CTIP2, CTIP1 is initially co-expressed with CTIP2, but its expression later becomes restricted to CPN and corticothalamic projection neurons, becomes excluded from SCPN, and is overall restricted to primary sensory areas. We find that the strategy of small molecule SIRT1 inhibition additionally promotes CTIP2^+^/CTIP1^-^ subtype distinction (**Figure 3B**). Based on both CTIP2^+^/SATB2^-^ and CTIP2^+^/CTIP1^-^ subtype distinction in the context of *Fezf2* expression, these data indicate that SIRT1 inhibition enhances and enables *Fezf2* refinement of neocortical subtype identity toward SCPN.

Given the known roles of SIRT1 in cortical neural progenitor differentiation and neuronal survival (Prozorovski *et al*., 2008; Li *et al*., 2008; Tiberi *et al*., 2012; Hisahara *et al*., 2008; Herskovits and Guarente, 2014; Cai et al., 2016; Iwata *et al*., 2020) and post-mitotic cortical neuron genomic stability (Dobbin *et al*. 2013), we next tested an alternative theoretical hypothesis that SIRT1 inhibition might potentially alter the proportions of progenitors and neurons, giving an impression of post-mitotic subtype distinction, while instead acting at the progenitor level. To the contrary, we find that the increase in proportion of CTIP2-expressing neurons is nearly completely compensated by the reduction of CTIP2^+^/SATB2^+^ dual expressing neurons (**Figure 3C,D**). Because the combined fraction of CTIP2- and SATB2-expressing neurons remains constant between samples, these experiments indicate that the subtype refinement phenotype is not due to changes in the proliferation of neocortical progenitors.

Although EX-527 and CHIC-35 are highly specific small molecule inhibitors of SIRT1 (Zhao *et al*., 2013), we pursued *Sirt1*-specific molecular knockdown with siRNA to further confirm whether SIRT1 is the main target of repression in primary dissociated neocortical neurons. We find that *Sirt1* knockdown in primary dissociated E12.5 neocortical neurons recapitulates the effect of small molecule inhibition of SIRT1, increasing both CTIP2^+^/SATB2^-^ and CTIP2^+^/CTIP1^-^ subtype-specific SCPN identity refinements (**Figure S4**).

### SIRT1 expression *in vivo* is CPN subtype-specific

We next investigated whether *in vivo* SIRT1 expression is consistent with the results of the screening approach in mES-derived neurons. We used immunocytochemistry to investigate SIRT1 protein localization in the developing mouse neocortex. At P4, we find that SIRT1 is expressed throughout the rostro-caudal extent of the neocortex, in layers II/III, V (at a relatively lower level), VI, and subplate (**Figure 4A**). While SIRT1 expression is broadly distributed, as previously reported (Hasegawa and Yoshikawa, 2008; Michan and Sinclair, 2007; Qin *et al*., 2006), we hypothesized that its level of expression varies in distinct neocortical subtypes. Strikingly, we find that SIRT1 expression is subtype-specific by E18.5, with near complete co-localization with SATB2-expressing CPN in layers II/III, V, and VI, and exclusion by CTIP2-expressing SCPN/CSN in layer V (**Figure 4B**). Similarly, at P4, SIRT1 expression is excluded from increasingly mature SCPN/CSN (**Figure 4C**).

**Figure 4.**
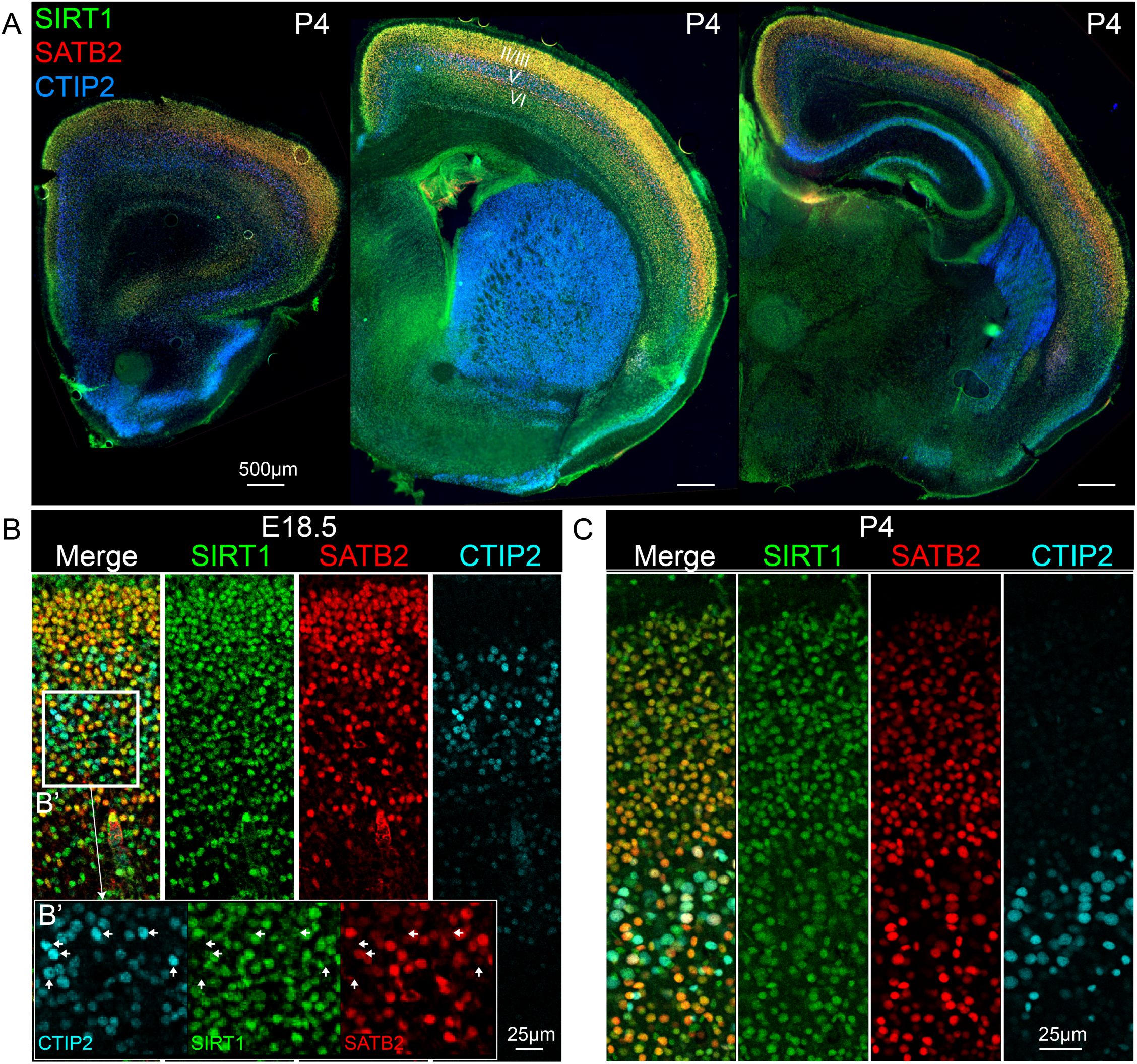
SIRT1 is differentially expressed by CPN and SCPN/CSN *in vivo*. (A) At P4, immunocytochemical labeling indicates that SIRT1 is expressed along the entire rostral-caudal extent of the neocortex, in layers II/III, VI, and subplate (50 μm coronal section, wide-field fluorescence imaging). (B) At E18.5, SIRT1 expression in deep layers of motor cortex is predominantly restricted to SATB2-expressing neurons, and is absent or expressed at quite low levels by CTIP2-expressing neurons (50 μm coronal section, confocal fluorescence imaging). (C) At P4, SIRT1 expression in deep layers of motor cortex is almost completely restricted to SATB2-expressing neurons (50 μm coronal section, confocal fluorescence imaging).

We next asked whether SIRT1 is differentially transcribed in pure populations of retrogradely-labeled CPN versus CSN, the important subtype of SCPN in layer V that project axons to the spinal cord. Using an existing microarray-based comparative gene expression analysis of retrogradely-labeled CPN and CSN (Arlotta *et al*., 2005; Molyneaux *et al*., 2009), we find that SIRT1 is the only differentially expressed histone deacetylase (HDAC) throughout post-mitotic neocortical differentiation at E18.5, P3, P6, and P14, with highest expression by CPN at all ages (**Figure S5**). Combined with the protein expression data in **Figure 4**, these results demonstrate that *Sirt1* mRNA and protein are specifically and highly expressed by SATB2-expressing CPN subtypes during corticogenesis, and are expressed at significantly lower levels by CTIP2-expressing SCPN/CSN during early, middle, and late stages of subtype identity refinement. These *in vivo* findings support the CPN-specific expression of SIRT1, and its relative exclusion from SCPN/CSN and other neocortical neurons.

Together, these data describe a context-specific role for SIRT1 in fine-tuning SCPN/CSN post-mitotic identity refinement during late embryonic neocortical development. Identified by high content, small molecule screening of epigenetic factors, SIRT1 inhibition enhances SCPN/CSN molecular refinement among primary dissociated E12.5 neocortical neurons and complements the approach of using transient *Fezf2* modRNA expression to promote SCPN/CSN identity refinement among heterogeneous mES-derived neocortical-like neurons.

## Discussion

Our data indicate that *Sirt1* inhibition or knockdown approaches are effective for the refinement of ES-derived SCPN/CSN *in vitro,* and suggest that SIRT1 is functionally important for the refinement of SCPN/CSN identity *in vivo* during mouse neocortical development. This is likely also relevant for human iPSC directed differentiation into SCPN/CSN. From the initial screening experiment **(Figure 2)**, we find that inhibition of SIRT1 preceding *Fezf2* induction enhances SCPN/CSN identity refinement in mES-derived neocortical neurons. Within primary mouse neocortical neurons, we identify that *Sirt1* inhibition, by either small molecule or knockdown approaches, promotes mature molecular refinement of *Fezf2*-mediated SCPN/CSN identity (**Figure 3**, **Figure S4**). Although *Sirt1*-null mice have not yet been assessed for subtype-specific deficits in the neocortex, their gross neocortical anatomy (*e.g.* intact corpus callosum, absence of Probst bundles) appears intact (Cheng *et al*., 2003; McBurney *et al*., 2003; Michán *et al*., 2010), suggesting that *Sirt1* is not required for CPN specification.

SIRT1 is a ubiquitously expressed NAD-dependent histone deacetylase (HDAC) with context-dependent roles in neocortical differentiation (Hisahara *et al*., 2008; Tiberi *et al*., 2012). At early developmental stages, SIRT1 regulates neurogenesis within neocortical progenitors by repressing the Notch-Hes pathway (Tiberi *et al*., 2012). Later in development, SIRT1 is ubiquitously expressed, with minimal enrichment in the upper layers of mouse neocortex at E14.5 and at 10 months of age, although the level of SIRT1 expression by specific neocortical subtypes had not been previously assessed (Hasegawa and Yoshikawa, 2008; Michan and Sinclair, 2007; Qin *et al*., 2006). We find that neocortical SCPN/CSN have markedly reduced SIRT1 expression in mid- to late-corticogenesis, in contrast to deep and superficial layer CPN that are relatively enriched for SIRT1 expression (**Figure 4**, **Figure S5**). Based on these results, SIRT1 should be considered to have neuronal subtype specificity as a chromatin modifier in the neocortex (Molyneaux *et al*., 2005; MacDonald and Roskams, 2008; Molyneaux et al., 2009).

Multiple lines of evidence indicate that SCPN/CSN identity refinement (*in vivo* and in mES-derived neurons) requires both *Fezf2* and a permissive molecular context during differentiation. First, transient *Fezf2* expression is sufficient to generate SCPN/CSN in *Fezf2*-null mice at E12.5, but not at E15.5 (**Figure 1**). At later ages (E13.5 through P7), a higher dose or duration of *Fezf2* expression (e.g., by plasmid vector) can re-specify alternate neocortical subtypes toward most aspects of SCPN/CSN identity (Molyneaux *et al*., 2005; Chen *et al*., 2008; Rouaux and Arlotta, 2013; De la Rossa *et al*., 2013). However, after E15.5, mis-expression of *Fezf2* does not induce Ctip2 expression in most neurons, highlighting context specificity (Chen *et al*., 2008; Rouaux and Arlotta, 2013; De la Rossa *et al*., 2013). Together, these prior *in vivo* findings indicate that E12.5 is the optimal molecular context to most completely enable *Fezf2*-mediated specification of SCPN, induction of Ctip2 expression, and stable epigenetic silencing of Satb2. Consistently, *in vitro*, high dose *Fezf2* induction (plasmid or viral mediated) by neocortical-like neurons does not alone induce SCPN/CSN identity (Wang *et al*., 2011; Sadegh, unpublished data, 2011). Moreover, when *Fezf2* modRNA is induced within mES-derived neocortical cells at a time approximating E12.5 neocortical differentiation, it alone does not significantly increase SCPN/CSN subtype-specific transcription factor expression (**Figure S2**). These *in vitro* data indicate that, although the timing of *Fezf2* expression is important, this cannot be accomplished by targeting cells with a suboptimal chromatin landscape, as would be expected in maturation-stalled ES-derived neocortical neurons (Sadegh and Macklis, 2014).

CPN are an evolutionarily more recent and diversified subtype of neocortical neurons, and likely employ multiple sequential epigenetic mechanisms in their specification, molecular refinement, and maturation (Molyneaux *et al*., 2009; MacDonald and Roskams, 2009; Kishi and Macklis, 2010; Fame *et al*., 2011; Fame et al., 2016a; Fame et al. 2016b). At late stages of maturation of layer 2/3 CPN (*e.g.* eight postnatal weeks in mice), the widely expressed methyl binding protein MeCP2 is required for proper development and/or maintenance of dendritic complexity and soma size (Kishi and Macklis, 2004, 2010; Kishi *et al*., 2016). At earlier stages of CPN development, SATB2 is required for proper differentiation, indirectly guiding chromatin remodeling by binding to matrix attachment regions (MAR) and recruiting HDAC enzymes through a binding partner, SKI (Britanova *et al*., 2005; Britanova *et al*., 2008; Alcamo *et al*., 2008; Gyorgy *et al*., 2008; Baranek *et al*., 2012). Together – with varying extents of CPN-specificity – SIRT1, SATB2/SKI, MeCP2, and likely others might coordinate chromatin remodeling in CPN at distinct stages of development.

The postmitotic subtype-specificity of SIRT1 expression in the neocortex is also remarkable because SIRT1 is implicated in the oxidative stress response and survival of neurons (Li *et al*., 2008; Prozorovski *et al*., 2008). Because SIRT1-expressing CPN might be resistant to metabolic insults, it raises the possibility that SCPN/CSN, by virtue of having reduced SIRT1 expression, might be more sensitive to metabolic stress. SCPN and especially the subpopulation of corticospinal neurons (CSN) are the brain neurons that selectively degenerate in amyotrophic lateral sclerosis (ALS; Ozdinler and Macklis, 2006; Zang and Cheema, 2002). Intriguingly, *Sirt1* over-expression has been shown to promote short-term survival of dissociated neocortical neurons mis-expressing ALS associated mutant SOD1 (Kim *et al*., 2007). More broadly, non-specific HDAC inhibitors show neuroprotective properties in mouse models of ALS (Petri *et al*., 2006; Rouaux *et al*., 2007).

Overall, our findings indicate subtype-specific functions for SIRT1 in the molecular refinement of neocortical SCPN/CSN versus CPN identity. Importantly, these results demonstrate the utility of combining epigenetic priming with subtype-specific transcription factor induction in ES (and not unlikely, human iPSC) directed differentiation. These results provide a proof-of-concept strategy for specific and progressive enhancement of directed CSN/SCPN or other subtype differentiation from pluripotent cells. These results further suggest that subtype-specific epigenetic modulation might enhance optimal *in vitro* generation of other diverse neocortical neuron subtypes.

## Methods

### RNA synthesis and transfection

Synthetic modified RNA (modRNA) was generated as previously described (Warren *et al*., 2010). Briefly, RNA was synthesized with the MEGAscript T7 kit (Ambion, Austin, TX). A custom ribonucleoside blend was used, comprising 6 mM 5’ cap analog (New England Biolabs), 7.5 mM adenosine triphosphate and 1.5 mM guanosine triphosphate (USB, Cleveland, OH), 7.5 mM 5-methylcytidine triphosphate and 7.5 mM pseudo-uridine triphosphate (TriLink Biotechnologies, San Diego, CA). Transfections of modRNA and multiple siRNA targeted against *Sirt1* and *Satb2* (both from Santa Cruz) were carried out with RNAiMAX (Invitrogen), as per the manufacturer’s instructions.

### Cell culture and differentiation

Feeder-free E14Tg2a (Baygenomics) mouse embryonic stem cells (mES) were passaged on gelatin-coated (0.1% gelatin, StemCell Technologies) cell culture treated plastic dishes using established media and cell culture techniques (Sadegh and Macklis, 2014). For differentiation, mES were plated at low density (5,000 cells/cm^2^) on gelatin-coated plastic dishes in ESC medium, and cultured as described (Gaspard *et al*., 2009). Cyclopamine (Calbiochem) was added from day 2 to day 10 in the differentiation medium at a final concentration of 1 uM. After 10 to 14 days of differentiation, cells were trypsinized, dissociated, and plated on poly-lysine/laminin (Becton-Dickinson) coated glass coverslips, and allowed to grow for 4–14 days in N2B27 medium (Gaspard *et al*., 2009).

### High-content small molecule screening

A high content screening protocol was adapted from Makhortova *et al*. (2011). Briefly, mES were seeded at 5,000 cells per well in 96-well plates, and treated in duplicate at 10 μM, 1 μM, and 0.1 μM with individual compounds from the screening library, a custom set of 80 chemicals affecting histone deacetylases, methyltransferases, and kinases. Most small molecules, including EX-527 (Sigma), CHIC-35 (Sigma), and nicotinamide (Sigma), were re-suspended in DMSO, according to the manufacturer’s instructions.

96-well plates were scanned by an automated confocal microscope (PerkinElmer Opera) at 20X magnification with separate fluorescence exposures from a UV light source and 488, 546, and 647nm lasers. Image analysis was performed using Columbus software (version 2.3.0; PerkinElmer; see also **Figure S3C,D**), which automatically set the boundaries of cell nuclei based on Hoechst staining. These boundaries were optimized by manual inspection to exclude nuclear fragments or adjacent double nuclei based on the total area and staining intensity of Hoechst-positive nuclei. Next, the intensity of antibody labeling for each distinct transcription factor in each nucleus was quantified. The threshold for positive antibody labeling was manually established individually for CTIP2, SATB2, and CTIP1, compared to baseline labeling without primary antibody (omission of primary controls). This threshold for labeling a nucleus positive for individual transcription factor expression was calibrated to approximately 20% of the maximum average pixel intensity observed in other nuclei for that transcription factor. Setting this threshold for positive labeling was necessary because populations of mES-derived neurons express a near-continuum of transcription factor labeling intensities, creating a large population of cells with simultaneous faint nuclear expression of multiple transcription factors, which could be interpreted as indicative of atypical transcriptional regulation (“confused” neurons). This is in striking contrast with the expression levels exhibited by *bona fide* populations of primary dissociated E15.5 mouse neocortical neurons, using the same labeling and analysis methods. These primary neurons typically display much more distinctly grouped average pixel intensities for each nucleus, with largely trimodal labeling that can be quantitatively delineated (negative, low expression, high expression).

### Mice

*Fezf2*-null mice were the generous gift of S. McConnell (Chen *et al*., 2005). Wild-type CD1 mice were used in all other experiments (Charles River Laboratories). The date of vaginal plug detection was designated E0.5, and the day of birth as P0. Mice used in these experiments were handled according to guidelines of the National Institutes of Health (NIH), and all procedures were conducted in accordance with Harvard University’s institutional guidelines.

### Immunocytochemistry

Immunocytochemistry and both wide-field and confocal imaging were performed as previously described (Sadegh and Macklis, 2014). Primary antibodies and dilutions used were: rat antibody to CTIP2 (1:500, Abcam); mouse antibody to SATB2 (1:200, Abcam); rabbit antibody to SIRT1 (1:250, Millipore); rabbit antibody to CTIP1 (1:500, Abcam); rabbit antibody to GFP (1:500, Invitrogen); chicken antibody to NESTIN (1:500, Novus Biologicals); mouse antibody to TuJ1 (1:500, Covance); mouse antibody to MAP2 (1:500, Sigma); and mouse antibody to NeuN (1:250, Millipore). Alexa fluorophore conjugated secondary antibodies from Invitrogen were used at a dilution of 1:1000. Hoechst 33342 counterstain was used to visualize nuclei (1:3,000, Invitrogen).

## Author contributions

C.S. and J.D.M. designed research; C.S. performed research; C.S. and J.D.M. analyzed data; C.S. and J.D.M. wrote the manuscript. W.E. provided expertise for the application of synthetic modified mRNA technology, reagent support, and discussion for this work. T.A., L.D., and L.L.R. provided expertise for small molecule screening, reagent support, and discussions.

## Acknowledgments

We thank Dr. Derrick J. Rossi for his laboratory’s technical and reagent support for modRNA generation, testing, and optimization. We thank A. Gee, Dr. K. Haston, and C. Goldstein (L. Rubin laboratory) for sharing their expertise in cell culture and high content screening, sharing of reagents, and technical assistance. We also thank Drs. T. Wuttke and H. Padmanabhan (J.D.M. laboratory) for help with primary neuron isolation, and expert advice, respectively.

This study was partially supported by a Harvard Stem Cell Institute (HSCI) program grant for the ES/iPS Directed Neuronal Subtype Differentiation collaborative program of the HSCI Nervous System Diseases Program, and by the Max and Anne Wien Professor of Life Sciences fund. This study was also supported by NIH grants NS45523 and NS41590, with additional infrastructure supported by NS49553 and NS75672, and with additional support from the Jane and Lee Seidman Fund for Central Nervous System Research, and the Emily and Robert Pearlstein Fund for Nervous System Repair (J.D.M.). C.S. was partially supported by T32 GM007753-30 (NIH Medical Scientist Training Program) and T32 AG000222-18 (Harvard Medical School, Division of Medical Sciences).

---

**Figure S1.**
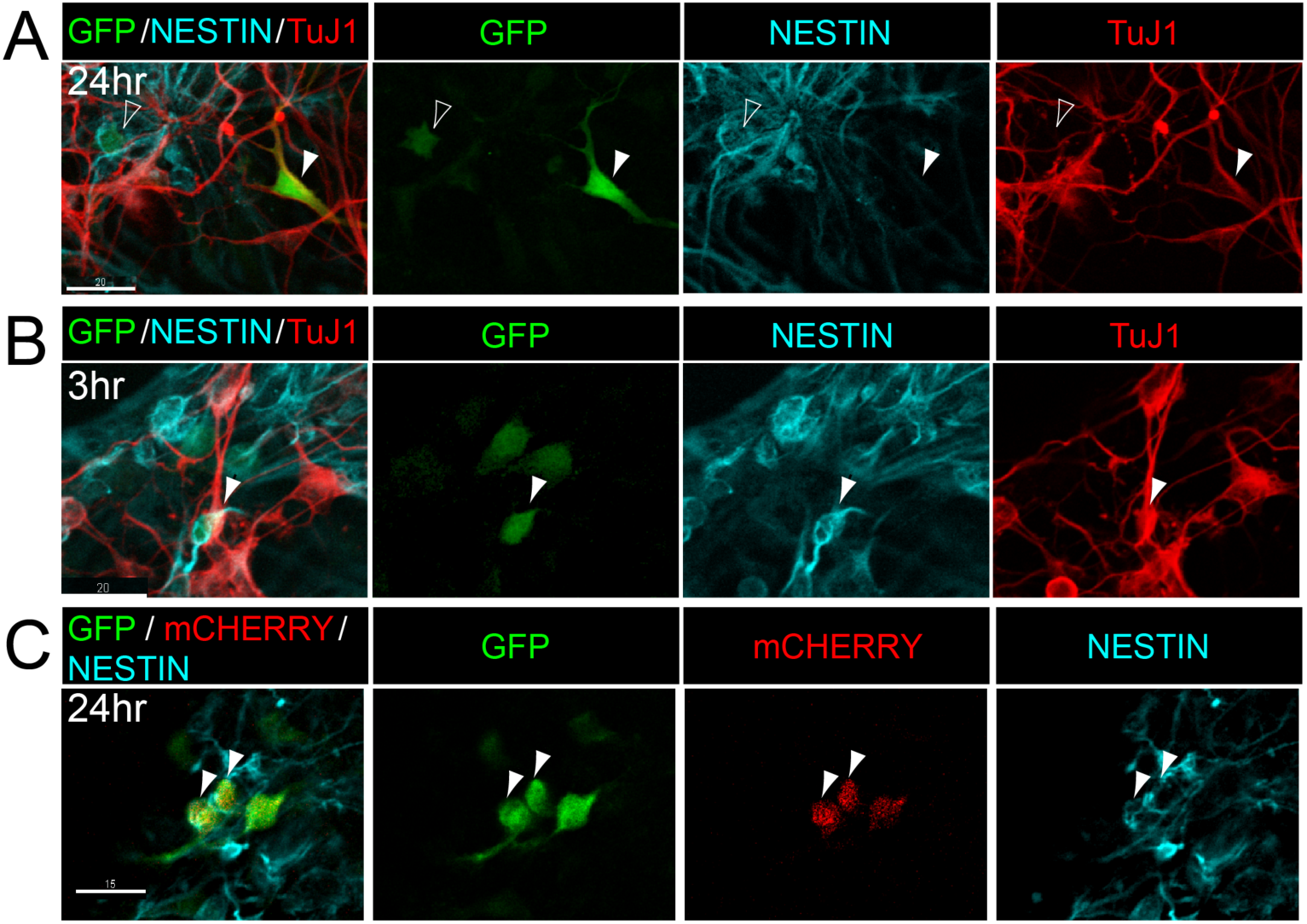
modRNA transfection is not biased to a specific neural population. (A) At 24 hours, *GFP* modRNA transfected mES-derived neocortical-like cells express GFP within progenitors (NESTIN-expressing, empty arrows), immature neurons (TuJ1-expressing, filled arrows), and other cells. (B) GFP is expressed by NESTIN-positive mES-derived cells as early as three hours following transfection with *GFP* modRNA. (C) *mCherry* and *GFP* modRNA are co-expressed by the same mES-derived cells 24 hours following transfection (native fluorescence).

**Figure S2.**
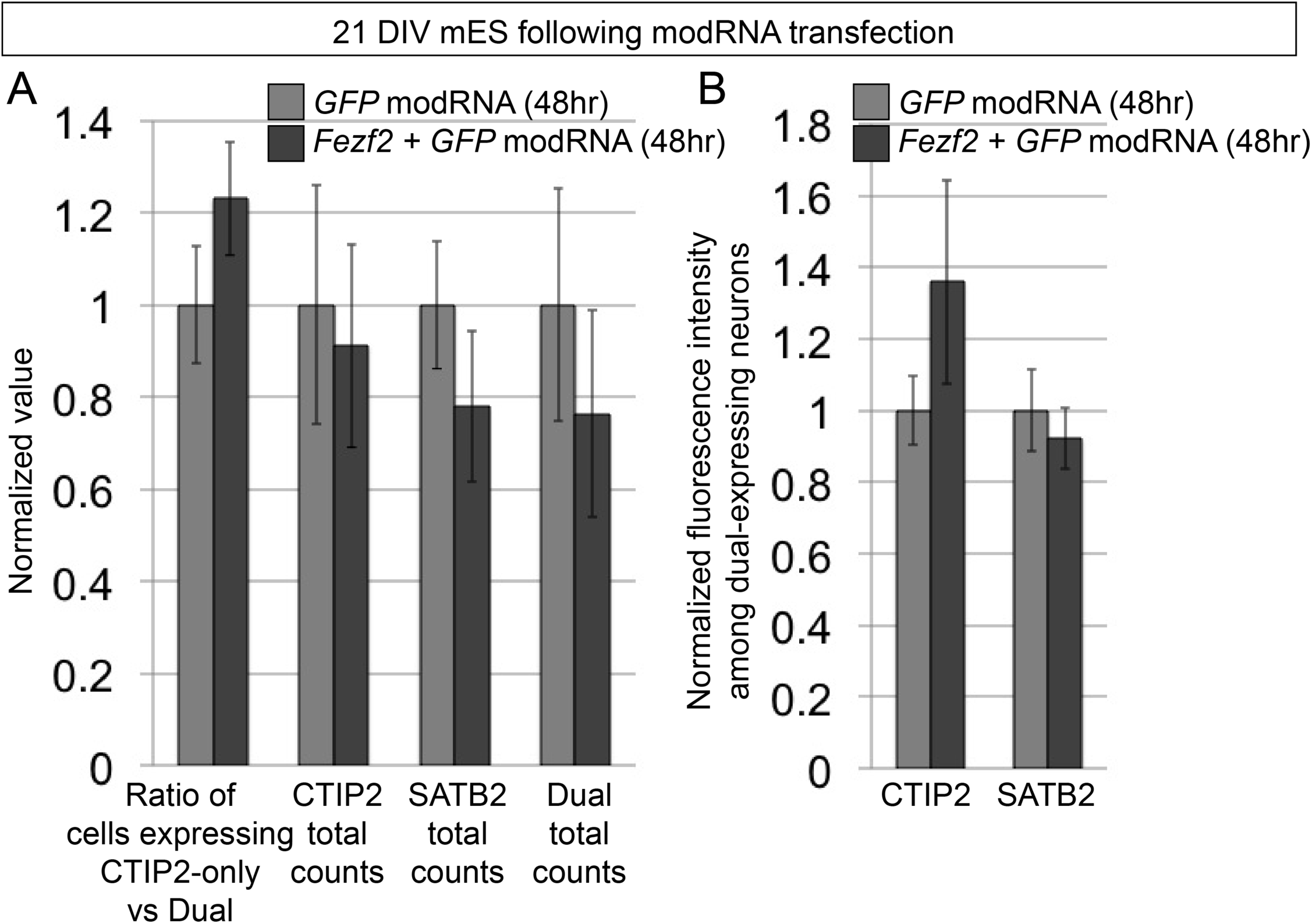
*Fezf2* induction in ES-derived neurons does not on its own significantly increase subtype distinction. (A) At 21 days in vitro (DIV), the ratio of CTIP2+/SATB2-neurons to CTIP2+/SATB2+ dual expressing neurons is not statistically significantly increased 48hrs after *Fezf2* and *GFP* modRNA co-transfection (dark grey) relative to *GFP* modRNA transfection alone (light gray) (though a trend suggests potentially modest increase of approximately 20%). The total number of CTIP2-expressing neurons is largely unaffected, as is the number of total SATB2-expressing and CTIP2+/SATB2+ dual expressing neurons (though the latter two display non-statistically significant trends toward decrease in number). (B) The intensity of CTIP2 expression within *Fezf2, GFP* modRNA co-transfected neurons increases relative to *GFP* modRNA controls. Data are presented as mean +/- s.e.m. (N=3; approximately 5,000 cells per condition, from 40 randomly sampled fields at 20x magnification).

**Figure S3.**
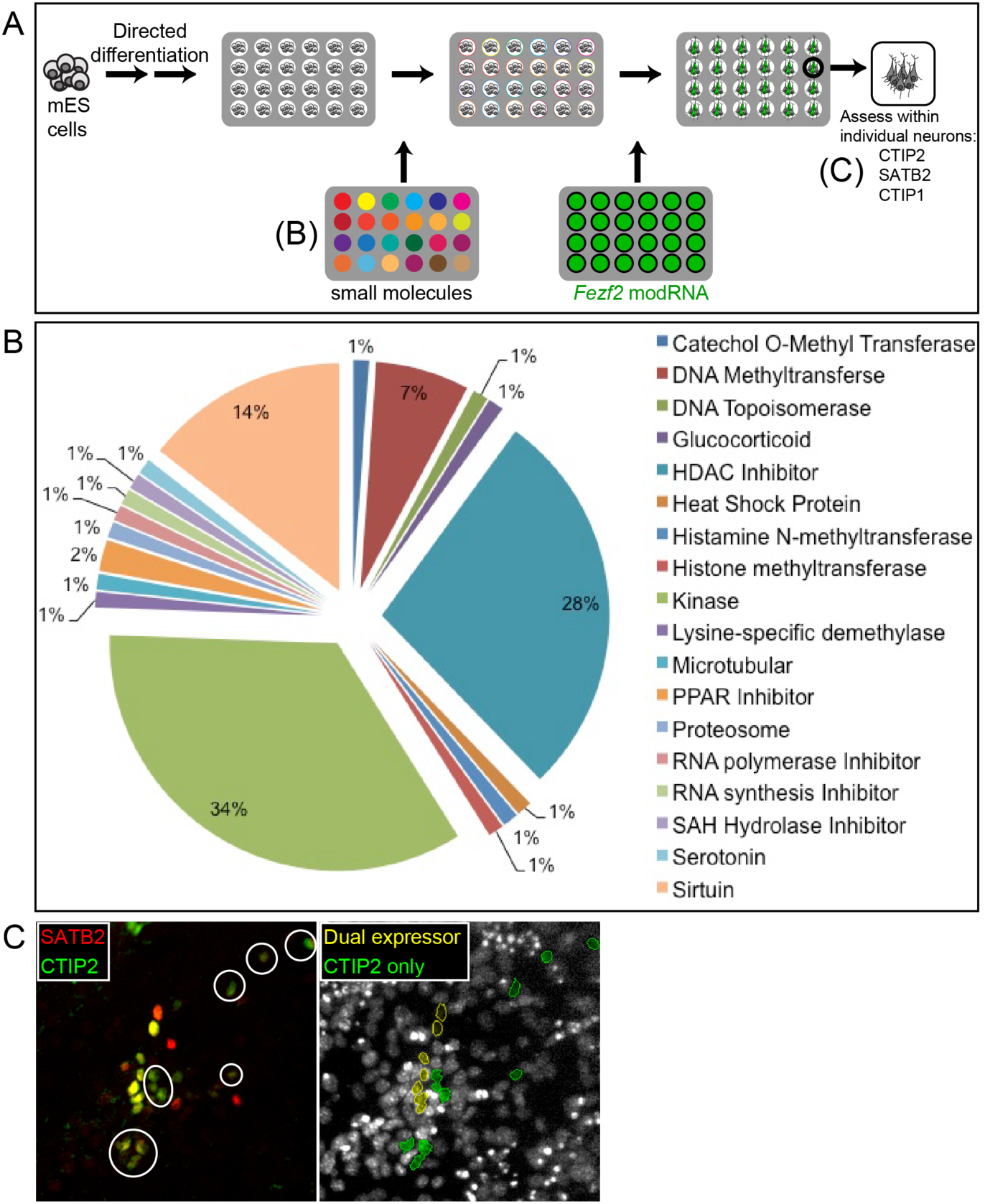
Design of high throughput small molecule screening protocol to identify potential regulators of subtype refinement within cortical-like neurons. (A) Schematic of screening strategy in 96-well plates. Monolayer mES differentiation to telencephalic progenitors is followed by the addition of a custom small molecule library; the composition of this library is described in (B). After small molecule incubation for six days, each well is transfected with *Fezf2* modRNA. Two days later, cells are fixed and immunolabeled for CTIP2, SATB2, and CTIP1. Automated imaging and fluorescence intensity thresholding algorithms distinguish and count neurons; an example of this is shown in (C). (B) The composition of a custom set of 80 chemicals regulating histone deacetylases, methyltransferases, and kinases is depicted in this pie chart. (C) Automated imaging and counting algorithms identify CTIP2 and SATB2 expression levels. Manually determined thresholds distinguish CTIP2+/SATB2-neurons (pseudo-colored green) from dual expressing CTIP2+/SATB2+ neurons (pseudo-colored yellow).

**Figure S4.**
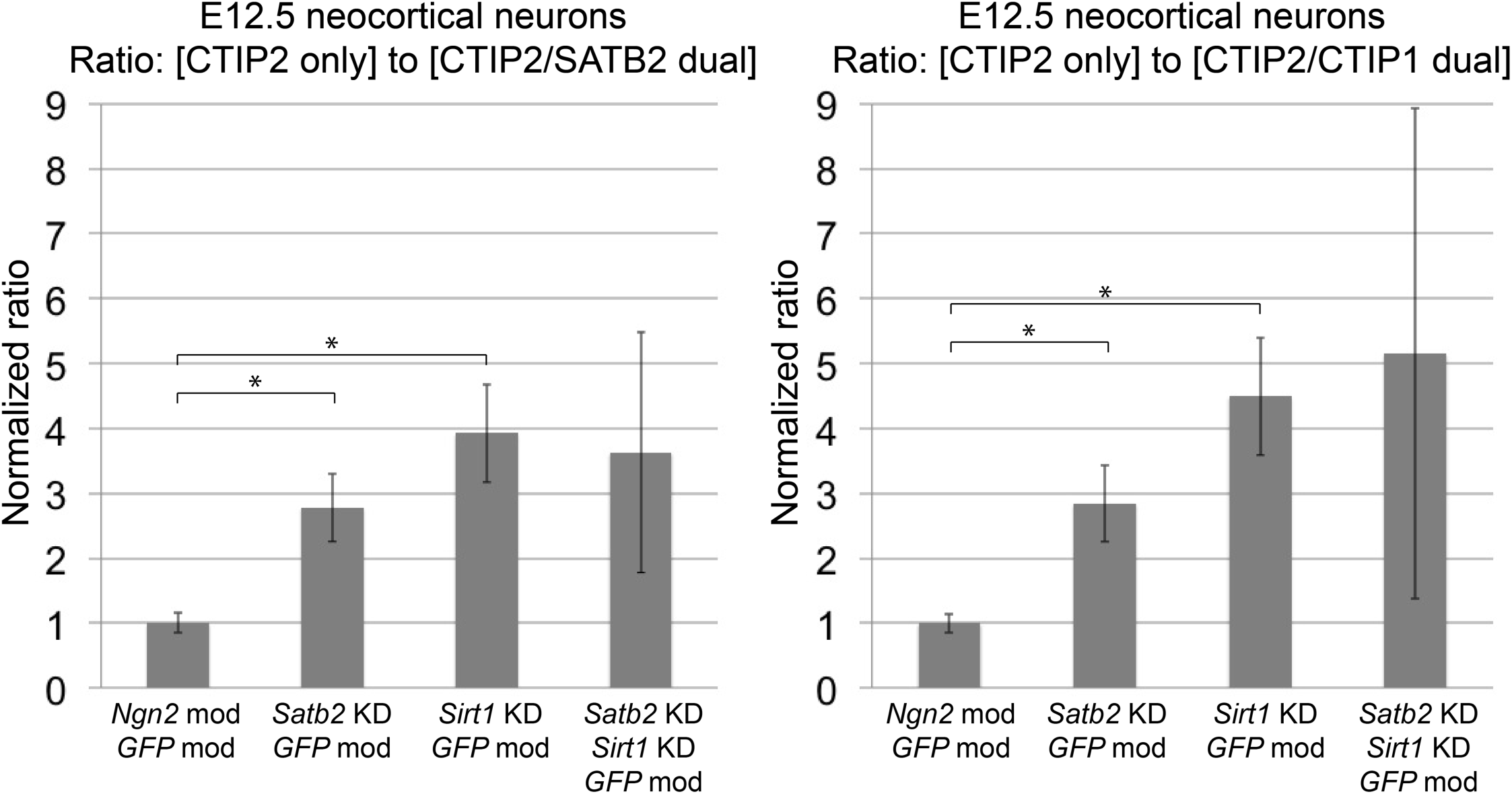
siRNA knockdown of *Sirt1* recapitulates the effect of small molecule inhibition of SIRT1. (A) CTIP2/SATB2 refinement increases with *Sirt1* knockdown (KD) in primary dissociated E12.5 neurons co-transfected with *GFP* modRNA, as compared to alternate conditions (co-transfection of *Ngn2* and *GFP* modRNA, or co-transfection of *Satb2* siRNA with *GFP* modRNA). (B) Consistent with multiple molecular refinements during neocortical projection neuron subtype distinction, CTIP2/CTIP1 refinement also increases with *Sirt1* knockdown. Data are presented as mean +/- s.e.m. (N=3; >10,000 nuclei screened per condition, from 60 randomly sampled fields at 20x magnification). *P < 0.05 (unpaired t-test).

**Figure S5.**
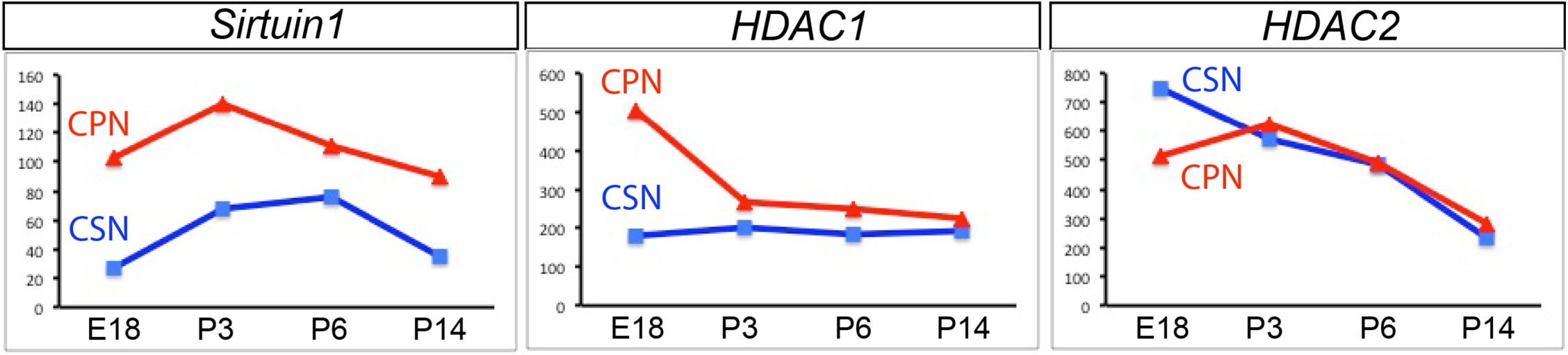
Sirt1 mRNA expression is CPN-specific in the neocortex at late embryonic and postnatal ages. (A) *Sirt1* mRNA expression is higher by CPN (red lines) than by CSN (blue lines) at E18.5, P3, P6, and P14; these populations were retrogradely labeled and purified by fluorescence-activated cell sorting (FACS) for comparative gene expression analysis at each time-point (data from Arlotta *et al*., 2005). (B) Other HDACs (*e.g.*, *HDAC1* and *HDAC2*) are not differentially expressed at all ages, using the same microarray data.

